# JACOBI4 software for multivariate analysis of biological data

**DOI:** 10.1101/803684

**Authors:** Denis Polunin, Irina Shtaiger, Vadim Efimov

**Affiliations:** Independent Researcher, Novosibirsk State University graduate, Novosibirsk, Russia; Institute of Cytology and Genetics SB RAS, Novosibirsk, Russia; Institute of Systematics and Ecology of Animals SB RAS, Novosibirsk, Russia; Novosibirsk State University, Novosibirsk, Russia; Tomsk State University, Tomsk, Russia

**Keywords:** genetic sequence, distance matrix, bootstrap, script

## Abstract

Biologists more and more have to deal with objects with non-numeric descriptions: texts (e.g. genetic sequences or even whole genomes), graphs, images, etc. There even could be no variables or descriptions at all when variability of objects is defined by similarity matrix. It is also possible to have too many variables (e.g. a magnitude of millions is reachable in mass spectrometry or genome research). In this case it is necessary to switch to object similarity matrices which drastically reduces dimensionality to hundreds or thousands. It is software developer’s responsibility to keep this use cases in mind and provide means for working with such data instead of shifting the problem to the users. Software should be more convenient for them and allow solving wider range of problems with fairly simple mathematical apparatus. In particular principal component analysis (PCA) is rather popular among biologists. But, the necessity of variables is an illusion. It’s enough to have a matrix of Euclidean distances between objects and apply method of the principal coordinates (PCo) (or multidimensional scaling for dissimilarity matrix, MDS) [1].

In the late 70s of the last century B. Efron proposed generating a set of new samples from the source sample EDF as a model for sample’s general distribution to get confidence estimation. He called it “bootstrap” [2]. For the statistical software developers this primarily means that PCo, MDS, and bootstrap should be implemented. Further, the use of bootstrap results in huge increase of repetitions of data analysis (from hundreds to millions of times) which is impossible to do in interactive mode. Therefore a part of the analysis requiring bootstrap should be written as a script in its entirety. Further user interaction should be eliminated. Obviously this process could be efficiently done in parallel.

There are multitude of tools for doing it varying from scripting languages like R or Python to specialized software packages like PAST, CANOCO, Chemostat, STATISTICA, and MATLAB. Researchers who are not versed in software development tend to use tools like PAST, even if they may not cover all their needs, including automating frequently performed tasks. However, automatic analysis is a key element for the upcoming era of bootstrap analysis.

We developed a simple and convenient package JACOBI4, which allows researchers without programming experience to automate multidimensional statistical analysis. Package and methods implemented in it can be useful in studies of both medical (gene expression for various diseases) and biological (regularities of molecular sequence variability) data. It goes without saying that the use of JACOBI4 is in no way limited to these examples. The package can be used directly, taking already developed scripts and editing them to fit own needs. Package JACOBI4 is freely available at [w1]. There are also articles available in which JACOBI4 is used to process real world data, as well as supplemental files containing JACOBI4 scripts and data for them.

## 1. Introduction

Research often requires multivariate data analysis. Usually, the following definition is implied: “Multivariate analysis is a set of methods used to analyze data that contain more than one variable” (https://www.camo.com/mva/). It is implicitly assumed that object descriptions are quantitative variables. However in modern age such understanding is evidently inadequate. Biologists more and more have to deal with objects with non-numeric descriptions: texts (e.g. genetic sequences or even whole genomes), graphs, images, etc. There even could be no variables or descriptions at all when variability of objects is defined by similarity matrix. It is also possible to have too many variables (e.g. a magnitude of hundreds of thousands or millions is reachable in mass spectrometry or genome research). In this case it is necessary to switch to object similarity matrices which drastically reduces dimensionality to hundreds or thousands. It is developer’s responsibility to keep this use cases in mind and provide means for working with such data instead of shifting the problem to the user.

Why multivariate analysis is necessary in such situations? The essence of multivariate analysis is geometric approach. Geometric approach means that objects are represented as points in multidimensional space and object similarity/dissimilarity could be seen as distance between points; preferably Euclidean distance. This model is more universal compared to conventional variable space. It is more convenient for biologists and allows solving wider range of problems with fairly simple mathematical apparatus. In particular principal component analysis (PCA) is rather popular among biologists. It operates on Euclidean variable space X reducing its dimensionality to form principal components − linear combinations of source variables. Thus, the illusion is created that a description of the variables is required. However there is another way.

More than half a century ago, J. Gower [1] discovered that if we calculate the matrix D of Euclidean distances between rows X (previously centered and normalized by columns), square this distances, double center and multiply by −1/2, then we get the XX^T^ matrix. Applying SVD to it: XX^T^ = PΛP^T^, we get the principal components U = PS (S = Λ^1/2^). For this reason J. Gower called this method the principal coordinates (PCo) analysis. However, it follows from the results of J. Gower that the matrix X itself is not needed and may not even exist in numerical form. To calculate the principal components of a certain set of objects, it is enough to have a matrix of Euclidean distances between them obtained by any way [3]. This is very useful in practice if the number of objects is significantly less than the number of traits that are becoming more common in biological research, especially molecular ones.

If we calculate the Euclidean matrix of distances between the rows of the matrix of PCs, then it will coincide with the initial matrix of Euclidean distances D. This property can be used to verify the calculations.

PCo is quite often used for dissimilarity matrices, for which it is unknown whether they are Euclidean distances between objects or not. In the case of non-Euclidean distances, some diagonal values of the matrix Λ will be negative. Small negative diagonal values can sometimes arise due to the accumulation of computational errors. All such “components”, as well as zero ones, should be excluded from consideration.

Instead of a distance matrix one can use a matrix of any coefficients of similarity/dissimilarity. In this case it is necessary to apply methods of multidimensional scaling (MDS). The results will always be the PCs in some Euclidean space [1]. (PCo has another name: metric multidimensional scaling with abbreviations MMDS or simply MDS. In our opinion, it is more correct to call MDS all methods of multidimensional scaling, but MMDS apply to PCo only).

One more thing for statistical software developers to keep in mind is the change in the ways of result confidence estimation. It is usually assumed the data is a sample from some general distribution. The parametric approach, which is becoming obsolete before our eyes, is based on the assumption that the distribution of data is known accurate to the parameters that are estimated from the sample. Moreover it is generally assumed the distribution to be normal (Gaussian). Therefore a normality test is always required. If the data do not meet this assumption, it is proposed to use nonparametric methods which typically are counterparts of parametric methods of confidence estimation, but on ranks only and distribution for ranks is known in advance. Such requirements are becoming less and less relevant especially in the Big Data era.

In modern approach the type of general distribution is extracted from the sample itself, since there is no other source to get it. In the late 70s of the last century B. Efron drew his attention to the fact that for any one-dimensional sample there always is empirical distribution function (EDF) [2]. According to Glivenko–Cantelli theorem with increasing number of objects the sample converges to its theoretical counterpart. B. Efron proposed generate a set of new samples from the source sample EDF and use this set as a model for sample’s general distribution to get required confidence estimation. He called it “bootstrap”. Further method development led to the idea of using smoothed EDF for bootstrap (it turned out to be a well-known statistics problem of restoring density of continuous and differentiable EDF). The method naturally was named “smoothed bootstrap” whereas the original became known as “naive bootstrap”.

For the statistical software developers this primarily means that both methods should be implemented. Further, the use of bootstrap results in huge increase of repetitions of data analysis (from hundreds to millions of times) which is impossible to do in interactive mode. Therefore a part of the analysis requiring bootstrap should be written as a script in its entirety. Further user interaction should be eliminated. The script execution is repeated to form statistic necessary to evaluate confidence. Obviously this process could be efficiently done in parallel.

There are multitude of tools for doing it varying from scripting languages like R or Python [4] to specialized software for statistics and analytics like PAST [5], CANOCO [w2], Chemostat [6], STATISTICA [w3], and MATLAB [w4]. Use of statistical packages is reviewed in Online Statistics Resources [w5]. Researchers who are not versed in software development tend to use tools like PAST, even if they may not cover all their needs, including automating frequently performed tasks. However, automatic analysis is a key element for the upcoming era of bootstrap analysis.

Our goal is to provide a simple yet powerful enough tool to allow researchers without programming background to automate statistical data analysis. In our opinion, this cannot be done in isolation from real biological tasks and data. Thus core functionality of the tool should rely on familiar to user concepts. To meet this goal we compiled the following list of things to keep in mind:

- Clarity of the module function.
- Data interoperability with external software.
- Consistent input data requirements for computational modules.
- Resilience to errors.

## 2. Architecture overview

JACOBI4 [w1] consists of a GUI for launching script execution and a set of modules performing operations on user data. Each module is an independent interchangeable component. Such separation is intended to increase stability of the tool and limit the impact of faulty module, allowing the user to work around the issue by using a different set of modules, e.g. PCA and PCo can be replaced with SVD and a few matrix operations.

## 3. Key features

### 3.1 Input data format

For the ease of use and interoperability with external software JACOBI4 modules operate on tabular data stored in CSV files. Even though the format is specified in RFC 4180, there are various implementations with idiosyncratic interpretations of it. To deal with this JACOBI4 has convention on the delimiter symbol and provides module for CSV format conversion. It’s expected for CSV files to have SEMICOLON separated values, and for TXT files to have TAB separated values.

Tabular data should have row and column keys, corner key and data matrix. Elements of the matrix could be numeric, string or empty (NaN). Modules have different restrictions on the input, e.g. multivariate analysis modules accept only tabular data with numeric values whereas some of the metric models accept empty values as well.

In typical workflow script operates on multiple files. It might contain merge, split, lookup and other algorithmic operations. A problem arises when input files have different encodings. There is no automated way of reliable encoding conversion so it’s up to the user to have it in sync.

### 3.2 Scripting language

For a user not versed in programming it could be a daunting task to write a script. It is expected the user would compose script from examples by altering them or copying pertinent blocks.

With the intention to simplify the matter we decided to use CSV files to store data processing scripts. This allows users to create, view and modify scripts with familiar tools like Microsoft Excel where it would be convenient for them to construct it.

The scripting language is a declarative one. Script consists of commands for processing data, variable declarations, loops and subroutines.

Script keywords declared in configuration file. It is possible to add synonyms or translations into different languages, e.g. variable assignment could be declared as “pi;=;3.14” or “pi;assign;3.14” or “pi;присвоить;3.14” or “pi; 分配;3.14”.

A module invocation consists of module name followed by a set of parameters each in its own cell. Before script execution each module invocation is verified to have the right amount of arguments and arguments having proper types.

Any script could be used as a subroutine with parameters passed as predefined variables “argument_0”, “argument_1” etc.

#### 3.2.1 Module description and internationalization

Command names and keywords could be translated into different languages. Since script stored as a table there are no restrictions on the symbols used in module names which also results in the ability to write script in somewhat natural language. JACOBI4 provides English, Russian and partially Chinese translations with the ability to extend or adjust it.

Modules information stored in accessible to user CSV files. Each line contains description of a new module with parameter type information, an alias to existing module or keyword and path to executable. It is possible for the user to add new module list file to provide convenient naming for the modules or add new translation.

The tricky part is to match encodings between user script and modules description files. We ended up looping over various encodings to find one that does not produce syntax and semantic errors. In the case when script contains errors there is no encoding information so we show errors for every encoding checked.

#### 3.2.2 Extensibility

There are two ways to extend functionality of JACOBI4. The first one is to write a subroutine in scripting language and add its description to module list file. This allows using this subroutine as if it is a module. It is expected for the user to accumulate gradually a library of their own convenient subroutines.

The second possibility is to add new module as a command line executable. It is expected this approach will rarely or never be used since our target audience are users without programming background.

### 3.3 Computation modules

#### Multivariate analysis

2B-PLS (Two-Block Partial Least Squares); PLS-regression (Partial Least Squares regression); SVD (Singular Value Decomposition); PCA (Principal Component Analysis); PCo (Principal Coordinate Analysis); NMDS (non-Metric Multi-Dimensional Scaling); MLR (Multiple Linear Regression); single linkage (Cluster Analysis); NNMF (Non-Negative Matrix Factorization).

#### Metrics

correlation; Euclidean metric; Minkowski metric; Manhattan distance; Hamming distance; Jaccard index; Jukes-Cantor distance; Kimura distance; Mantel test; rank Mantel test; p-distance; spectrum distance.

#### Matrix operations

matrix product; Fisher angular transformation; Fisher z-transformation; centering; double centering; distribution function; Fisher transformation; matrix elementwise operations; elementwise operations with two matrices; length normalization; max normalization; quantile normalization; sigma normalization; square normalization; sum normalization; statistics: mean, standard deviation, variance, skewness, excess kurtosis, sum, sum of squares, min, max; make binary matrix; transpose.

#### Resampling

bootstrap; permutation.

#### Matrix preparation

matrix splitting; append rows; append columns; append rows with header; convert group vector to matrix; convert line to matrix; convert matrix to line; copy columns; copy rows; delete rows by column values; delete rows with not number values; exchange columns by key value; exchange rows by key value; extract value; initialize matrix; insert value; insert column labels; insert diagonal; insert row labels; label count; label replacer; merge matrices; shift matrix values; move rows up; replace values by regular expression; replace cell values by values from another file; replace columns by ranks; replace if contains; replace labels; sort by column; sort by row; split by column; split by row; split columns by substring; split rows by substring; trim matrices.

#### File operations

convert CSV (allows to change separator used in CSV file); copy file; create file; delete file.

#### Other

echo; encoding conversion; plot; print header; subroutine.

## 4. Practical use of JACOBI4

JACOBI4 is mainly intended for calculations and does not yet have its own graphics. For illustrative purposes, users need to use other software.

### 4.1. Comparison with other tools

Our goal is in providing a simple yet powerful enough statistical tool to a researcher, e.g. biologist or physician without programming (as well as physical and mathematical or engineering) background. For such user tool should be as simple and convenient to use as possible, implying it should be free of charge, have support of batch data processing, have simple yet universal data structure with easy, understandable, intuitive, preferably close to natural, input language without unnecessary restrictions and complications.

JACOBI4 designed specifically for this kind of users. This “ecological niche” is currently empty.

Such well known tools as SAS, STATISTICA, SPSS, MATLAB and many more are commercial; for this reason they are excluded from our comparison. The most popular non-commercial tools among biologists are R and PAST. Despite its 20+ years of development, PAST unfortunately does not have batch data processing. Whereas it is an essential feature for various reasons. Firstly, it is required to be able to verify computation correctness. With scripting, it is always possible to get back to the text and debug it. In interactive mode, it is very easy to make a mistake and miss it. Secondly, well-tried script could be used as many times as needed. Particularly, it allows the user to create and reuse blocks of well-tried data processing code where all that left to do is to replace input/output file names and tweak a few parameters. It is also possible to repeat blocks of code in a loop, but they should be self-contained, i.e. they should not require user interaction. It is especially important for the bootstrap computations.

R is non-commercial, scriptable and universal at the same time. It can do everything. By no means it is a coincidence, R dominates biological data processing domain. However it is designed for the advanced user whereas the majority of biologists and physicians, in our experience, are not advanced and they don’t use R. There are multiple reasons for this. Firstly, R is too complicated for such users due to their lack of appropriate education. Secondly, they are very busy on their primary job; despite all benefits they just don’t have time to master R. Such users have their own, close and understandable biological tasks. The necessity to digress from it to dive into an area of unfamiliar and incomprehensible mathematical and programming structures and abstractions causes psychological discomfort. For example, R has rather complicated data model. It is clear that this is done for versatility, but this creates unnecessary problems for users. The R-package still maintains its archaic convention of using abbreviations and shortened names of functions. This is understandable to programmers but why should the user adapt to this? For the ordinary user function’s purpose should be clear at a glance. The necessity of studying its documentation scares him off especially considering the sheer amount of functions in R and the many parameters. We do not browse every article on the Internet, but filter them by title. In JACOBI4 this issue is solved by using additional CSV file with mapping from user visible name to name of executable module. This file is accessible to user to add convenient for him module names. It makes scripts easy to comprehend. More over such approach simplifies multilingualism problem. Module name can be represented as an arbitrary character sequence. The use of English, Russian and simplified Chinese module names has already been proven to work; it is also possible to mix module names from different languages in the same script.

### 4.2. How JACOBI4 can be used with an example dataset

We address two examples using JACOBI4 to process real data sets. Examples are specially selected by various to demonstrate the methods similarity and capabilities of the package.

#### 4.2.1 Gene expression in Huntington disease

Article [7] references gene expression dataset GSE3790-GPL96 for post mortem brain tissue comparison by three brain regions between patients with Huntington disease (HD) and controls. Data are taken from [d1]. Gene expression measured by GeneChip Affymetrix Human Genome U133A Array [HG-U133A].

Two Excel spreadsheet files are generated from the GSE3790-GPL96 array. The__GSE3790A.csv file contains 22283 probesets for gene expression profiles in 201 samples. The__GSE3790A-Inf.csv file contains, among others, quantitative (Age, Genotype1, Genotype2) and categorical (Health [cnt, dis] Grade [cnt, 0, 1, 2, 3, 4], Region [CN, FC, CB], Gender [F, M]) traits of each sample.

In each table, the first row consists of a corner key and column keys, the first column consists of the same corner key and row keys. In__GSE3790A.csv, columns correspond to samples; in__GSE3790A-Inf.csv, rows correspond. Any table in JACOBI4 can always be transposed with a single directive (command line).

The numbers of rows and columns of *data* in JACOBI4 scripts starts at $1, $0 is reserved for the *keys*.

To process HD data using JACOBI4 (current version is JACOBI_4.3.13), the script__Script_HD_A.csv was applied (the lines are automatically numbered in Excel and are provided only for ease of perception):

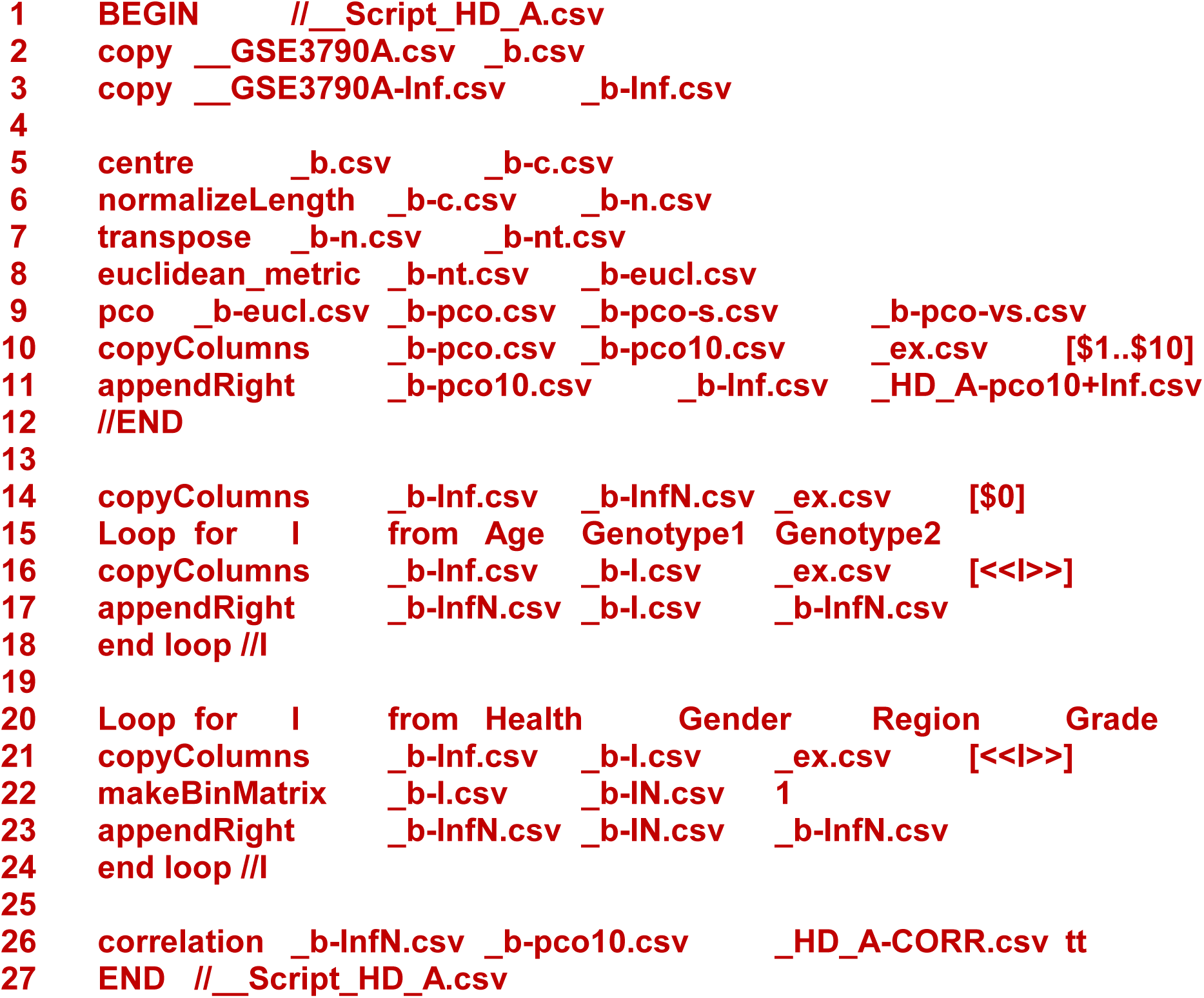

The commands in lines 5..11 standardize (center and normalize) the data table row by row (probesets), transpose table, calculate the Euclidean matrix of distances between rows (these are already samples), calculate the PCs by the Principal Coordinate method (PCoA), leave the first 10 PCs and append all the traits of the samples to the right. The resulting table is already suitable for graphing. But one more step is required for numerical analysis.

The commands in lines 14..18 in the loop form a table of quantitative traits for the samples. The commands in lines 20..24 in the loop add categorical traits in binary representation to it on the right. The binary representation is generated by the command on line 22. The command on line 26 calculates the correlation matrix (Table 1). Hereinafter, we did not consider correlation coefficients less than 0.3 due to small effective size.

**Table 1.** Correlation coefficients (×1000) between the PCs and the quantitative and binary traits of the samples

| REF | PC1 | PC2 | PC3 | PC4 | PC5 | PC6 | PC7 | PC8 | PC9 | PC10 |
| --- | --- | --- | --- | --- | --- | --- | --- | --- | --- | --- |
| Age | 14 | 206 | 12 | 38 | -32 | -212 | -86 | 21 | -174 | 60 |
| Genotype1 | 12 | 122 | -21 | 75 | -151 | -6 | 44 | 60 | -34 | 203 |
| Genotype2 | 40 | 242 | -143 | 172 | **-359** | -207 | 26 | -72 | 73 | 176 |
| Health_HDcnt | -41 | -249 | 156 | -179 | **337** | 234 | -12 | 38 | -49 | -193 |
| Health_HDdis | 41 | 249 | -156 | 179 | **-337** | -234 | 12 | -38 | 49 | 193 |
| Gender_F | 50 | 18 | 17 | 11 | -10 | -112 | -8 | 142 | -105 | -90 |
| Gender_M | -50 | -18 | -17 | -11 | 10 | 112 | 8 | -142 | 105 | 90 |
| Region_CB | **978** | 140 | -26 | 17 | 50 | 68 | 34 | 8 | -6 | -37 |
| Region_CN | **-615** | **587** | -149 | **-321** | **320** | 60 | -84 | -34 | 34 | -53 |
| Region_FC | **-356** | **-738** | 177 | **310** | **-376** | -130 | 51 | 26 | -29 | 91 |
| Grade_HD0 | -3 | 36 | 106 | 25 | -46 | -18 | 75 | 35 | -41 | 148 |
| Grade_HD1 | 19 | -14 | 0 | -51 | -9 | -82 | -44 | 162 | -56 | -12 |
| Grade_HD2 | 7 | 259 | -60 | 139 | -227 | -280 | -51 | -136 | 33 | 21 |
| Grade_HD3 | -19 | 67 | -262 | 55 | -184 | 64 | 142 | -171 | 30 | 37 |
| Grade_HD4 | 73 | -15 | -48 | 153 | -69 | 98 | -27 | 44 | 162 | 248 |
| Grade_HDcnt | -41 | -249 | 156 | -179 | **337** | 234 | -12 | 38 | -49 | -193 |
| $\lambda$, % | 17.4 | 13.0 | 5.9 | 3.3 | 2.7 | 2.3 | 1.6 | 1.3 | 1.3 | 1.2 |
| $\Sigma$, % | 17.4 | 30.4 | 36.3 | 39.6 | 42.3 | 44.6 | 46.3 | 47.6 | 48.9 | 50.1 |

Note: coefficients $\geq 0.306$ are highlighted (for $N=201$ p-value = $1.0 \times 10^{-5}$ ).

**Fig. 1.**
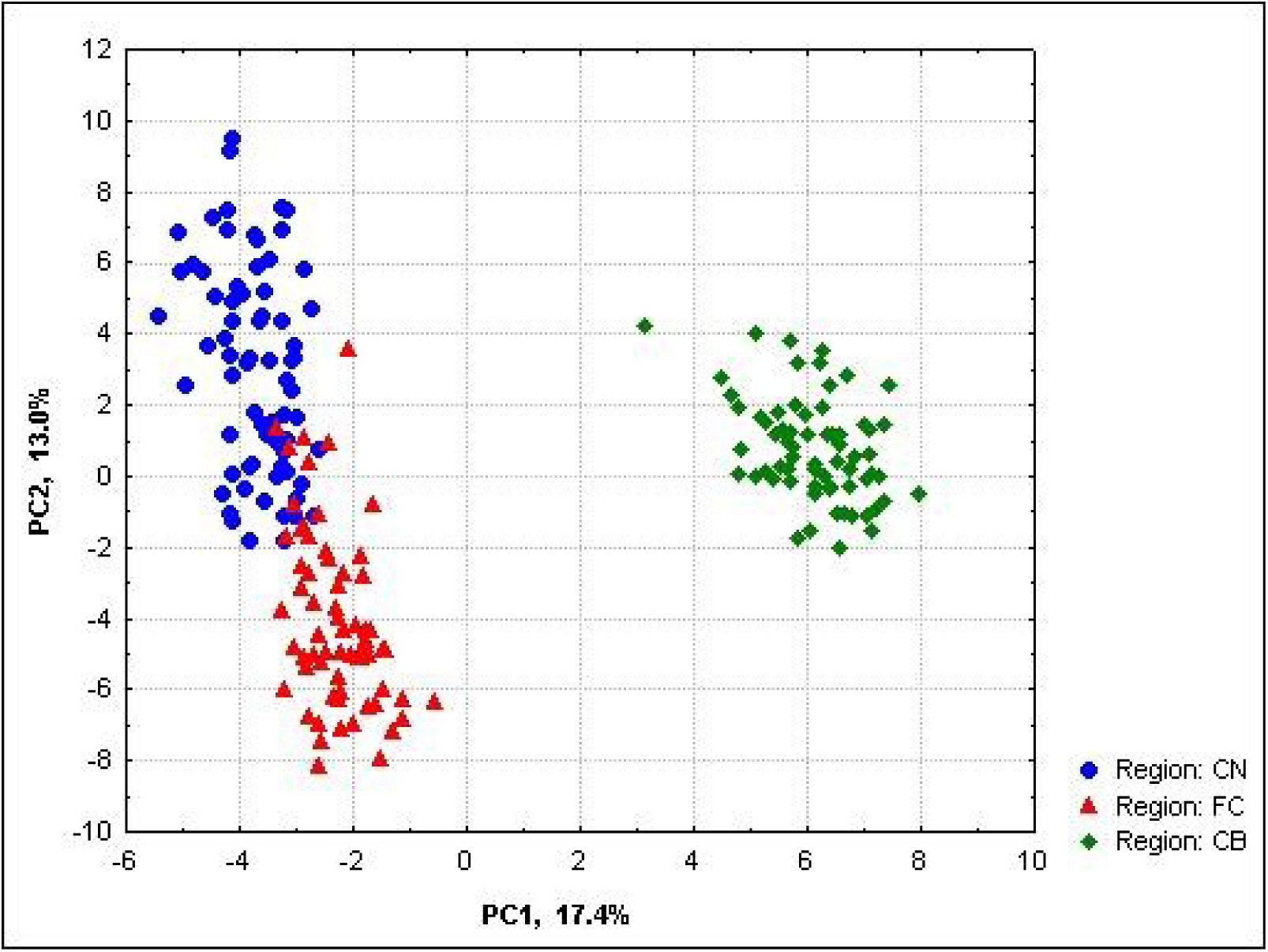
Configuration of brain tissue samples in the plane of the first two PCs of gene expression variability (CN – caudate nucleus, FC – frontal cortex, CB – cerebellum).

It follows from the Table 1 that the variability of the first two PCs is associated with differences in gene expression between brain regions (Fig. 1). Therefore, gene expression should be considered for each region separately. For this, some changes have been made to the script. Lines 2, 3, 20 are replaced by the following:

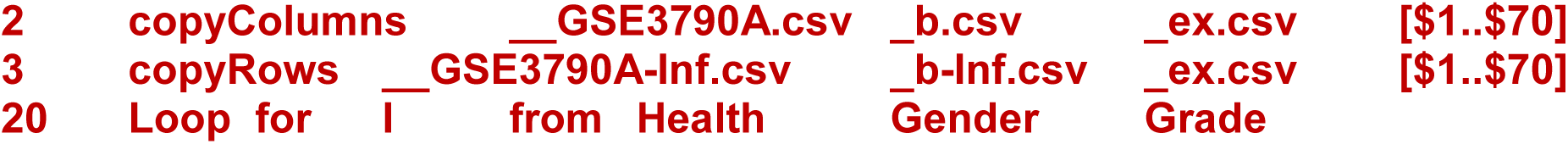

Commands 2, 3 leave in the table only information related to the region of the CN brain (caudatus nucleus, first 70 numbers). Command 20 removed categorical trait Region because it now consist only one value (CN). The results obtained after the script changes are shown in Table 2.

It follows from the Table 2 that the variability of gene expression between samples in the PC1 is associated with different degrees of HD manifestations, and from the Fig. 2 – that the degree of manifestation increases along the PC1.

It should be noted that information on quantitative (Age, Genotype1, Genotype2) and categorical (Health, Grade, Gender) traits of the samples was not used at all in the PCs calculations. It is needed only for their interpretation. PCs were calculated only by the multidimensional variability of gene expression profiles of samples from the CN brain region. The fact that PC1 correlates specifically with HD means that the maximum variation in gene expression within the CN region is associated with HD. Since it is PCoA that catches it, PC1 can be used to search of candidate genes, for example, through its correlations with probeset expression profiles. Note that a search in all regions at once may be unsuccessful, since during the processing it turned out that the variability of gene expression primarily depends on the brain region and only secondly on the degree of manifestation of HD.

An exception is the chaotic arrangement of samples with a zero degree of HD0, i.e. with the absence of clinical manifestations of the HD. But we need to keep in mind that the numbers of CAG repeats (Genotype 2) in these patients are in the range of 40–43, and this means (for living ones) a genetic predisposition to HD and, almost always, the clinical manifestation of this disease in the future.

Apparently, it is reasonable not to include such patients in either the HD class or the control class, but allocate them in a separate class of genetically predisposed to HD, but without the clinical manifestations of this disease [8].

The configuration of the samples in the plane is close to the obtained by us in [9]. The samples are the same, but the chip [HG-U133B] and the dataset (GSE3790-GPL97) are different [d1]. It goes without saying that one can compare the results with JACOBI4. Calculations can be repeated for other brain regions using small script replacements.

One can also calculate the correlations of all probesets with PC1 and select gene-candidates with the maximum modulus of correlation. The next logical step should be to build a prototype of the HD gene network from them. However, all this does not fit into the scope of this work.

**Table 2.** Correlation coefficients (×1000) between the PCs and the quantitative and binary traits of the samples from the CN region of the brain (caudatus nucleus)

| REF | PC1 | PC2 | PC3 | PC4 | PC5 | PC6 | PC7 | PC8 | PC9 | PC10 |
| --- | --- | --- | --- | --- | --- | --- | --- | --- | --- | --- |
| Age | 145 | 135 | 209 | 88 | -48 | 159 | 285 | 182 | 90 | 252 |
| Genotype1 | 247 | -62 | 140 | 51 | -68 | 108 | -77 | 12 | -127 | -123 |
| Genotype2 | **755** | -38 | 226 | -114 | -9 | 11 | -36 | -174 | -101 | -135 |
| Health_HDcnt | **-746** | 49 | -262 | 148 | 54 | -66 | 65 | 120 | 105 | 62 |
| Health_HDdis | **746** | -49 | 262 | -148 | -54 | 66 | -65 | -120 | -105 | -62 |
| Gender_F | 58 | 120 | 160 | 102 | -22 | 83 | -10 | -36 | -163 | 83 |
| Gender_M | -58 | -120 | -160 | -102 | 22 | -83 | 10 | 36 | 163 | -83 |
| Grade_HD0 | 86 | 238 | 70 | 20 | 87 | 116 | 21 | 7 | -217 | 116 |
| Grade_HD1 | 29 | -6 | 113 | -120 | -227 | 135 | -228 | -12 | -152 | 54 |
| Grade_HD2 | **556** | 19 | 227 | -82 | -13 | -7 | 67 | -26 | 154 | -149 |
| Grade_HD3 | **364** | -251 | -33 | 71 | 166 | -268 | 127 | -87 | -79 | -104 |
| Grade_HD4 | 134 | -115 | -120 | -128 | 55 | 243 | -77 | -193 | 51 | 113 |
| Grade_HDcnt | **-746** | 49 | -262 | 148 | 54 | -66 | 65 | 120 | 105 | 62 |
| $\lambda$, % | 13.9 | 10.9 | 6.1 | 5.1 | 3.3 | 2.5 | 2.2 | 2.1 | 2.0 | 1.9 |
| $\Sigma$, % | 13.9 | 24.8 | 30.9 | 36.0 | 39.3 | 41.8 | 44.0 | 46.1 | 48.2 | 50.0 |

Note: coefficients $\geq 0.306$ are highlighted (for $N=70$ p-value = **0.01**).

**Fig. 2.**
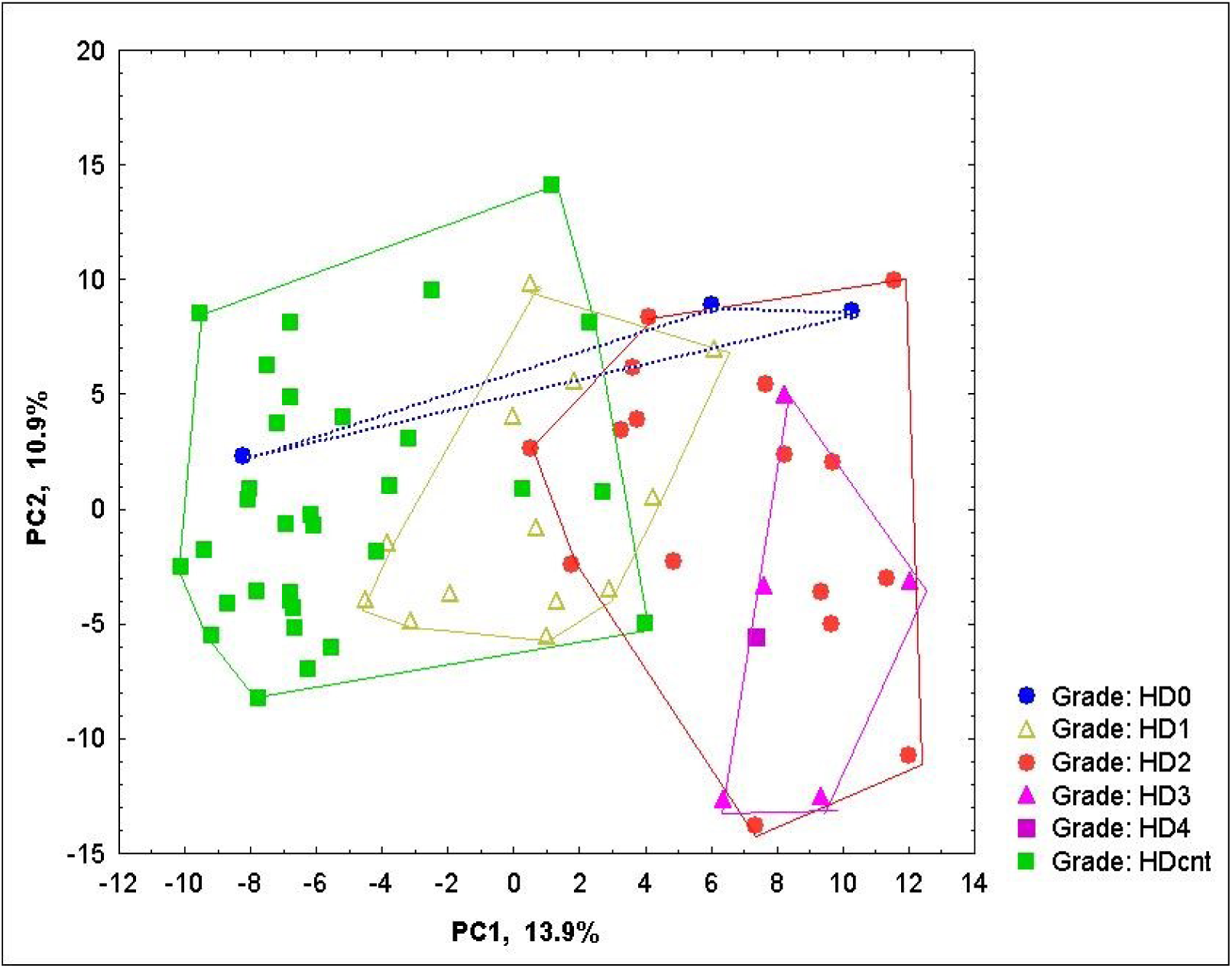
Configuration of tissue samples from the CN region of the brain (caudatus nucleus) in the plane of the first two PCs of gene expression variability.

#### 4.2.2. Principal Component Analysis for *Cytb* sequence (PCA-Seq)

The article [3] provides an example of the processing of *Homo sapiens Cytb* protein sequence by the PCA-Seq method. Now we adress a sequence from a species belonging to a different taxon and having slightly different genetic code. *Drosophila melanogaster* (Fruit fly) *Cytb* sequence P18935 is taken from [d2]. It consists of 378 amino acids (AAs).

The function of *Cytb* protein in all animals is to transfer energy through the mitochondrial membrane. *Cytb* crosses it in both directions several times. It is required to reveal patterns of *Cytb* sequence and, after that, to associate them with known transmembrane topology features such as AA location in α-helix and/or membrane.

To do this, the sequence was processed using the SS_SPIDER2 algorithm [10] and the Quick2D set from the MPI Bioinformatics Toolkit [11] which contains eight algorithms (SS_PSIPRED [12], SS_SPIDER3 [13], SS_PSSPRED4 [14], SS_DEEPCNF [15], SS_NETSURFP2 [16], TM_TMHMM [17], TM_PHOBIUS [18], TM_POLYPHOBIUS) [19]). Prefix SS denotes that algorithm predicts AA location in α-helix, TM − in membrane.

The__CYB_DROME.csv file contains 378 rows. Each row consists of a serial number, a one-letter AA code, and binary codes of all nine algorithms results (1, if AA is located in α-helix/membrane; 0, otherwise). The supplementary file__AcidCod.csv contains one-and three-letter abbreviations of AAs.

To process data using JACOBI4 (current version is JACOBI_4.3.13), the script__Script_AcidDomens.csv was applied:

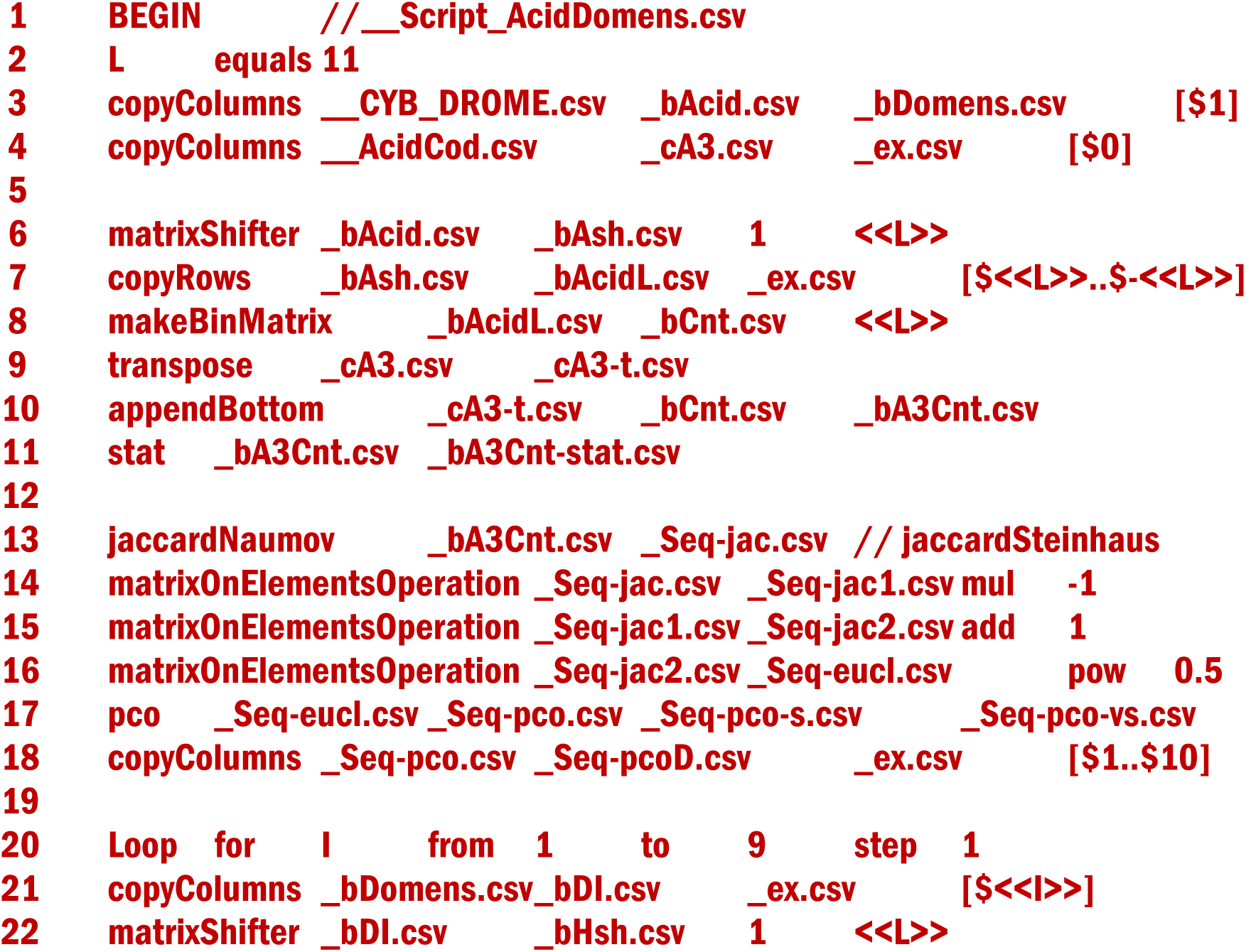
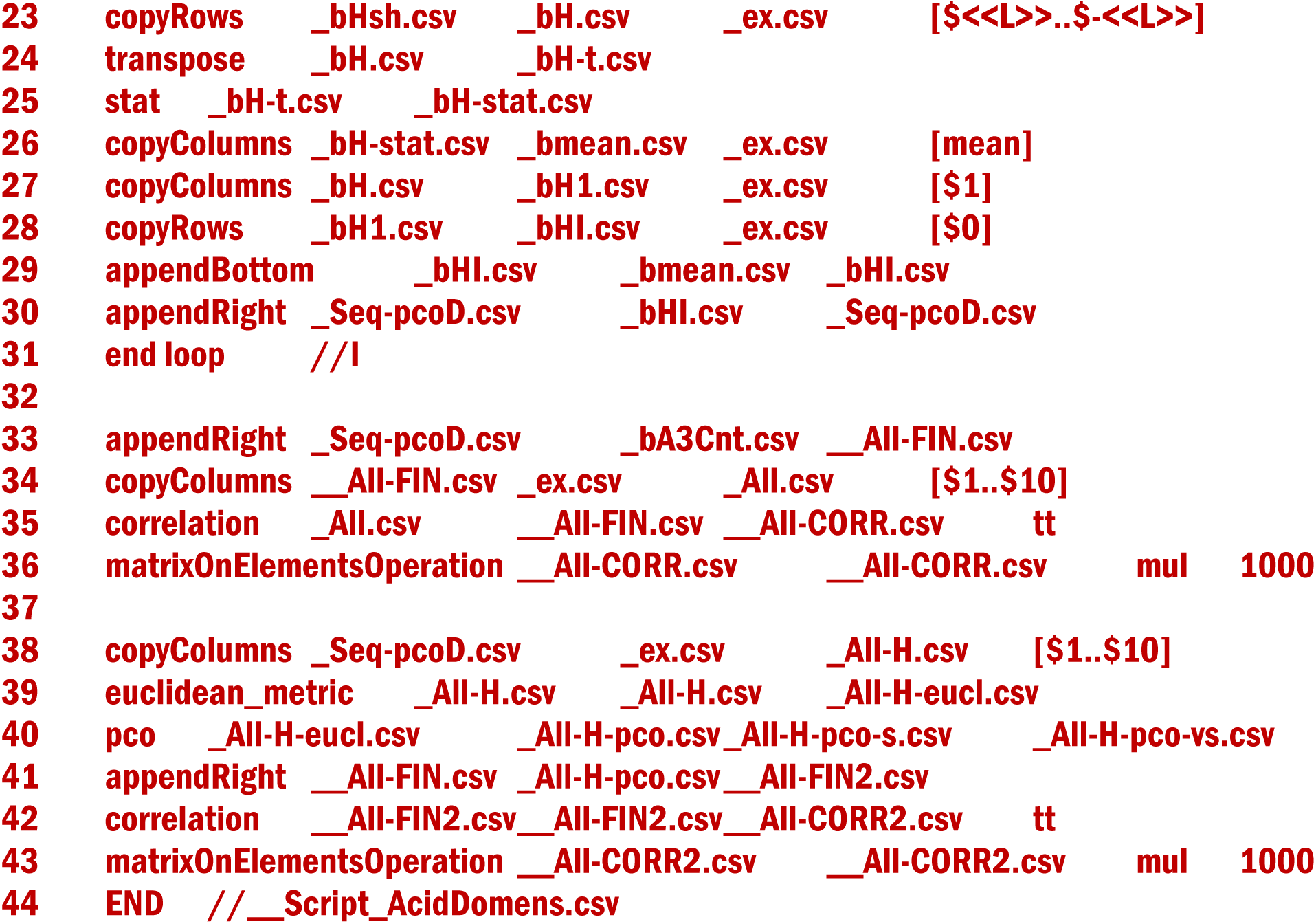

In line 2 of the script, the length of the fragment L is specified, namely L = 11. It is known that the length of one turn in an α-helix is ~ 3.6..3.7 AAs. Three turns (10.8..11.1) approximately correspond to the 11 AAs.

The command in line 3 splits the__CYB_DROME.csv file into a one-letter AA column and nine columns of the binary codes of the AA location in α-helix/ membrane. Command 4 prepares a column vector of the three-letter amino-acid codes [ala,..., tyr].

Command 6 duplicates the original column L-1 times, shifting each down by one element relative to the previous. Command 7 leaves only full rows in the array. Thus the table “fragment×shift” size of 368×11 is formed. Command 8 calculates for each row the number of occurrence of each AA in fragment, forming a new table. In lines 9..10, the title of the new table is drawn up.

Commands 13..16 calculate the Euclidean matrix of distances between fragments by transforming the Jacquard−Steinhaus distance [20, 21]. The following commands 17..18 calculate the PCs(AA) from the obtained distance matrix using the PCoA and leave the first 10 PCs in the same way as in 4.2.1.1.

Commands 20..31 calculate, in a cycle for nine binary traits, a fraction of each fragment location in α-helix/membrane and add it as new trait (with prefix SS or TM) to the table of the PCs.

Command 33 adds a table with the occurrences of all AAs in fragment. Commands 34..36 calculates the correlation matrix between the PCs and the traits of the fragments and multiplies it by 1000.

Command 38 separates a table consist SS_ and TM_traits only. Command 39 calculates the Euclidean matrix of distances between fragments. Command 40 calculates new PCs (SS+TM) from the obtained distance matrix using the PCoA.

Command 41 adds a table with PCs (SS+TM). Commands 42..43 calculates the correlation matrix between PCs (AA) and the all traits of the fragments including PCs (SS+TM) and multiplies it by 1000 for the convenience of printed reproduction and visual perception (Table 3).

The calculation results are shown in Table 3 and in Fig. 3.

Why is the Euclidean distance calculated using the Jacquard coefficient of similarity in line 13..16? First of all, in order to show that there are such opportunities. Calculations according to ordinary Euclidean distance are equivalent to ordinary PCA. However, it is not necessary to be limited only by them. Calculations through the Jacquard coefficient are becoming increasingly popular and often lead to more interesting results.

From table 3, it is seen that the protein sequence fragments location in α-helix/ membrane is mainly reflected by PC1(AA) (**7.2%** of the total variance). Results of all nine algorithms correlate precisely with it, and with sufficiently large correlation coefficients (**0.565..0.755**). Three rather weak coefficients (**0.328..0.337**) corresponds to the PC4(AA) only. PC1(SS+TM) correlates best of all with PC1(AA) (**0.761**) obtained from the results of all nine algorithms for predicting the secondary structure of the sequence (**75.9%** of the total variance). PC2(SS+TM) (**11.8%)** reflects the difference in the results of prediction of AA location in α-helix either membrane (its correlation coefficients with all SS variables are positive,with all TM variables are negative). However, this PC, as well as PC3(SS+TM) (**4.5%**), does not practically correlate with any of the PCs(AA).

From table 3, it is seen that the protein sequence fragments location in α-helix/ membrane is mainly reflected by PC1(AA) (**7.2%** of the total variance). Results of all nine algorithms correlate precisely with it, and with sufficiently large correlation coefficients (**0.565..0.755**). Three rather weak coefficients (**0.328..0.337**) corresponds to the PC4(AA) only. PC1(SS+TM) correlates best of all with PC1(AA) (**0.761**) obtained from the results of all nine algorithms for predicting the secondary structure of the sequence (**75.9%** of the total variance). PC2(SS+TM) (**11.8%)** reflects the difference in the results of prediction of AA location in α-helix either membrane (its correlation coefficients with all SS variables are positive,with all TM variables are negative). However, this PC, as well as PC3(SS+TM) (**4.5%**), does not practically correlate with any of the PCs(AA).

It should be noted that any information or assumptions about the secondary structure of the sequence or AA features were not used in the calculations of the Pcs(AA) at all. They were calculated by the similarity of the sequence fragments between themselves according to the set and amount of AAs, i.e. by the primary sequence of AAs only.

Nevertheless, PC1(AA) and PC1(SS+TM) correspond to each other very well, despite the fact that PC1(AA) reflects 7.2% only of the variability of the sequence fragments by the set and quantity of AAs (Fig. 3).

**Table 3.**
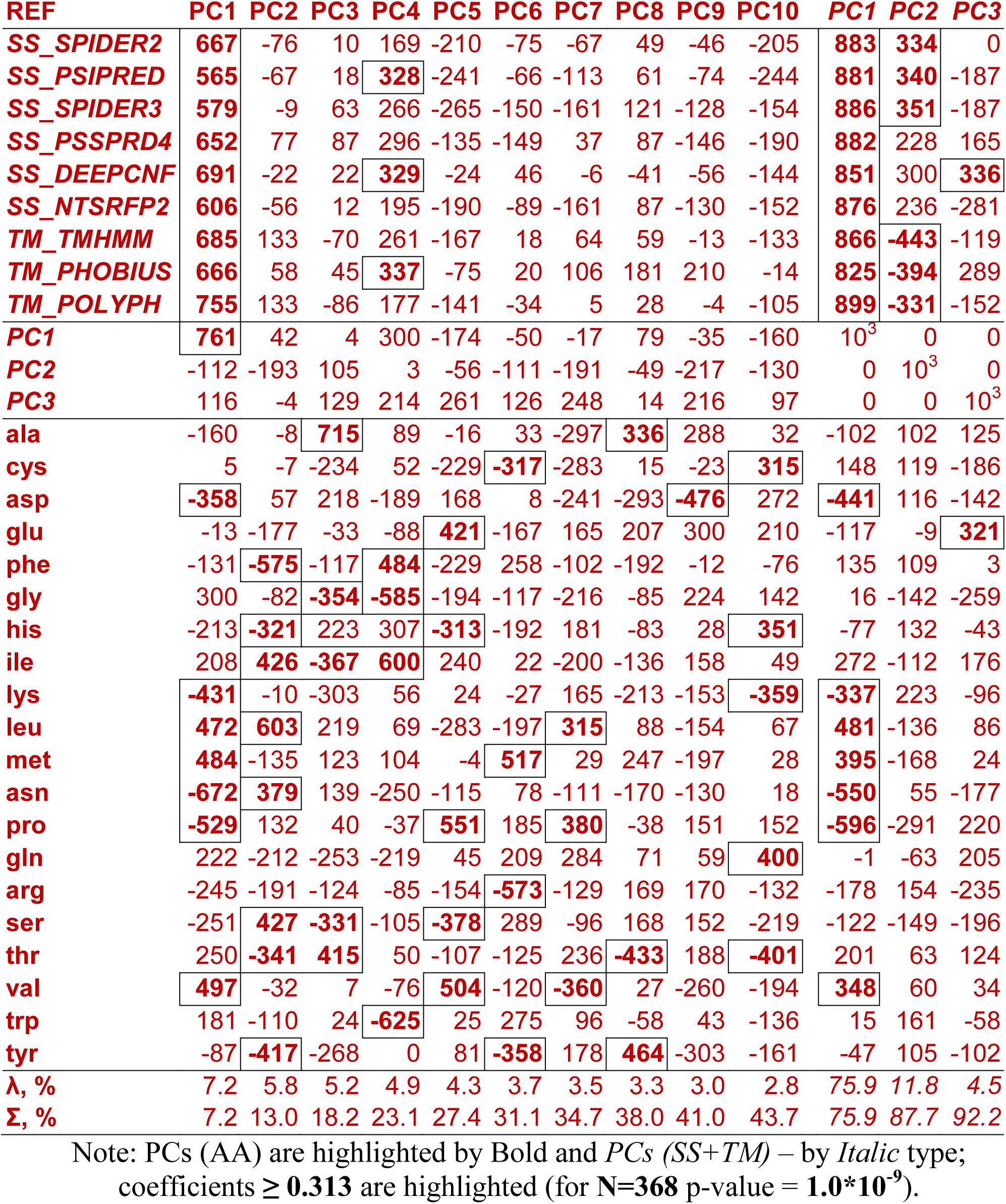
Correlation coefficients (×1000) between the PCs (AA and *SS+TM*) and quantitative traits of the fragments of the *D. melanogaster Cytb* sequence P18935

Since all PCs(AA) are orthogonal to each other, other PCs(AA) can add some more information about the secondary structure (possibly PC4(AA)). In addition, they certainly carry information about other sequence patterns that do not yet fall into the researchers view area. We believe this is a topic for future research.

**Fig. 3.**
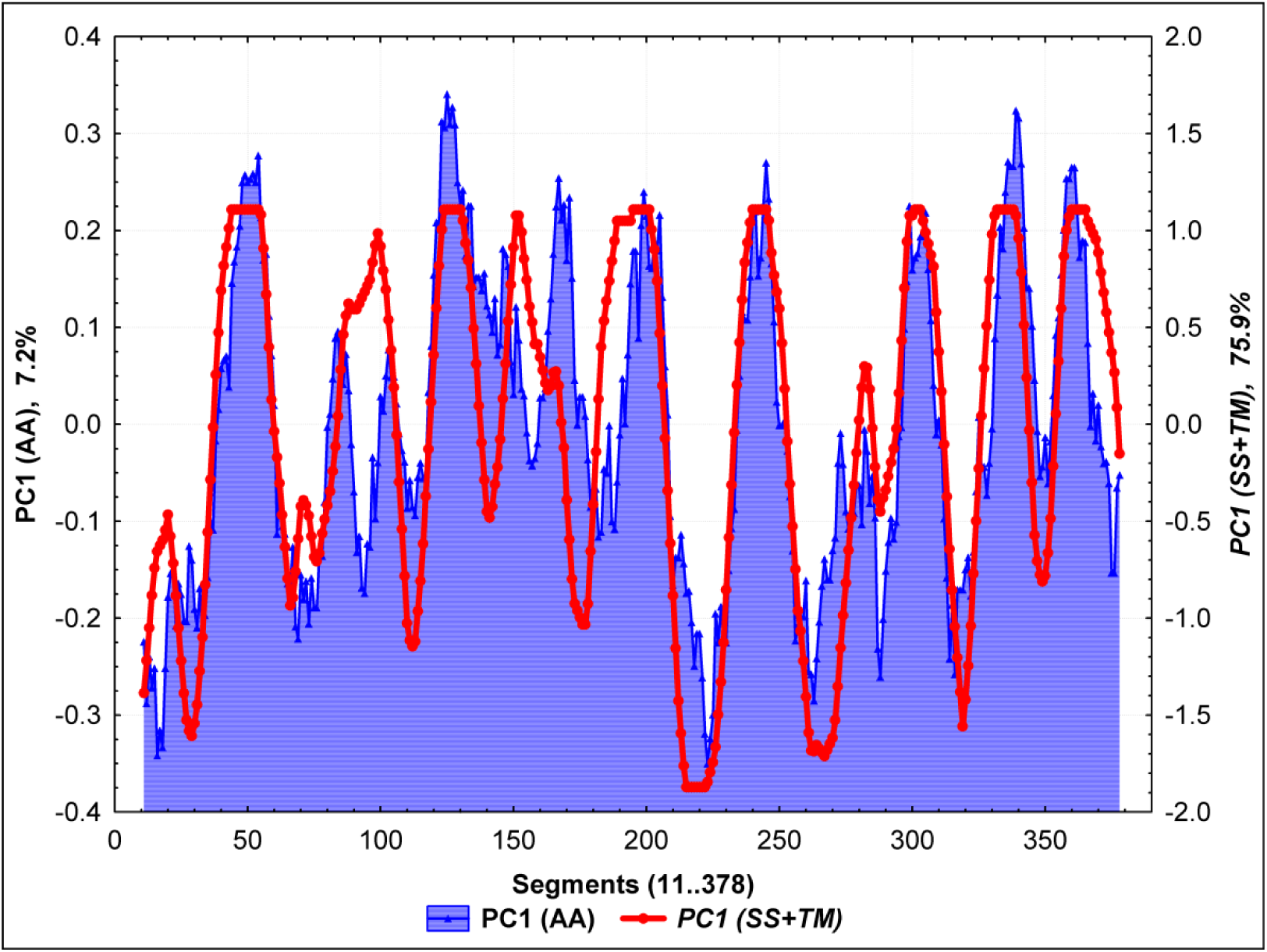
Dynamic of the **PC1(AA)** and ***PC1(SS+TM)*** of the sequence fragments variability (*D. melanogaster Cytb* P18935) (r= **0.761**; N= **368**; p**< 10^−12^**).

Thus, PCA-Sec and its implementation through JACOBI4 allow us to obtain potentially useful information on the protein secondary structure quickly and directly from the primary sequence without any assumptions.

## Conclusion

It follows from the above examples that package JACOBI4 and the methods implemented in it can be useful in solving both medical (gene expression for various diseases) and biological (patterns of the molecular sequences variability) problems. It goes without saying that the scope of the package is by no means limited to these examples. The package can be used directly, taking already developed scripts and editing them to fit own needs. We plan to expand the library of scripts.

Package JACOBI4 is freely available at [w1]. There are also articles available [3, 22] in which JACOBI4 is used to process real biological data, as well as supplemental files containing JACOBI4 scripts and data for them.

## Acknowledgments

The authors are grateful to Russian Federal Science & Technology Program of development of genetic technologies. This work was supported by the RFBR [#19-07-00658-a] and Budget Project [#0324-2019-0040].

